# Molecular characterization of inversion breakpoints in the *Drosophila nasuta* species group

**DOI:** 10.1101/2021.06.01.446624

**Authors:** Dat Mai, Doris Bachtrog

## Abstract

Chromosomal inversions are fundamental drivers of genome evolution. In the *Drosophila* genus, inversions have been widely characterized cytologically, and play an important role in local adaptation. Here, we characterize chromosomal inversions in the *Drosophila nasuta* species group using chromosome-level, reference-quality assemblies of seven species and subspecies in this clade. Reconstruction of ancestral karyotypes allowed us to infer the order in which the 22 identified inversions occurred along the phylogeny. We found a higher rate of inversions on the X chromosome, and heterogeneity in the rate of accumulation across the phylogeny. We molecularly characterize the breakpoints of six autosomal inversions, and found that repeated sequences are associated with inversion breakpoints in four of these inversions, suggesting that ectopic recombination is an important mechanism in generating inversion. Characterization of inversions in this species group provides a foundation for future population genetic and functional studies in this recently diverged species group.

## Introduction

Inversion polymorphisms have been studied extensively in *Drosophila* genetics since their first discovery over a century ago (Sturtevant 1917). Chromosomal inversions were first identified as suppressors of recombination in *Drosophila melanogaster* (Sturtevant 1917), and characterized subsequently in detail as structural alterations in polytene chromosomes across the *Drosophila* genus (Krimbas and Powell 1992).

Over the past century, chromosomal inversions have been recognized as a ubiquitous evolutionary phenomenon. Inversions are present in virtually all species and can have wide-ranging evolutionary effects. Inversions can help maintain coadapted gene complexes, reduce gene flow in hybrid zones, or restrict recombination between diverging sex chromosomes (Hoffmann and Rieseberg 2008). In addition to modifying the recombination landscape along a chromosome, inversions can also directly alter the structure or expression of genes found near inversion breakpoints (Calvete et al. 2012; Guillén and Ruiz 2012).

Despite being ubiquitous in nature and their putative widespread consequences, the evolutionary forces maintaining inversions are typically poorly understood. Several lines of evidence suggest that many inversions found in Drosophila and other species are adaptive. In particular, inversions often show seasonal, altitudinal and/or latitudinal clines, and polymorphic inversions are often associated with fitness-related traits (Hoffmann and Rieseberg 2008; Hoffmann, Sgrò, and Weeks 2004).

Genome-wide alignments between species allow us not only to detect the presence of chromosomal inversions but also to identify and characterize inversion breakpoint regions (Feuk et al. 2005; Ranz et al. 2007). Breakpoint sequences may shed light on the causes generating the inversion as well as on the functional consequences that the inversion might have had.

Here we identify chromosomal inversions and characterize their breakpoints in the *D. nasuta* subgroup. This species group contains about a dozen species that are distributed across South-East Asia. The karyotype of *D. nasuta* species consists of the X (Muller A), a large metacentric autosome (chromosome 2; Muller B, E) and a large acrocentric autosome (chromosome 3; Muller C, D), and the small dot chromosome. In *D. albomicans*, chromosome 3 fused to both the X and the Y chromosome, forming a neo-sex chromosome. Inversion polymorphism of the *nasuta* species group has been studied using cytogenetic techniques (Casu 1990; Lambert 1982;Pope 1987), and species in this group were found to be highly polymorphic for chromosomal inversions. However, no systematic characterization of inversions at the molecular level exists. Here we take advantage of high-quality chromosome-level genome assemblies for molecular characterization of inversions in the *D. nasuta* subgroup.

## Results

### Karyotype evolution in the *D. nasuta* subgroup

Ancestral linkage groups are conserved across the Drosophila genus and termed Muller elements (Muller 1940). Most flies in the *D. nasuta* subgroup have a conserved karyotype with a telocentric X (Muller A), a large metacentric autosome (Muller B and Muller E) and a telocentric autosome (Muller C and Muller D), and the small dot chromosome (Muller F). In *D. albomicans*, the telocentric autosome fused to both the ancestral X and Y, forming a neo-X and neo-Y chromosome. Our high-quality assemblies recovered each chromosome arm as a single contig (Mai, Nalley, and Bachtrog 2020). **Figure 1** gives an overview of global syntenic relationships across the species investigated, based on the location of protein-coding genes; genes are assigned to Muller elements (and color-coded accordingly). Consistent with previous studies within the Drosophila genus, syntenic comparisons on all Muller elements reveal a rich history of intrachromosomal reshuffling of genes (Bhutkar et al. 2008; **Figure 1**). Interestingly, while Muller C and Muller D genes are mixed up along the telocentric chromosome 3, no shuffling of Muller B and Muller E genes occurred on the metacentric chromosome 2. This suggests that paracentric inversions are more frequent in this group than pericentric inversions, consistent with observations in other Drosophila groups (Sperlich and Pfriem 1986; Krimbas and Powell 1992).

**Figure 1.**
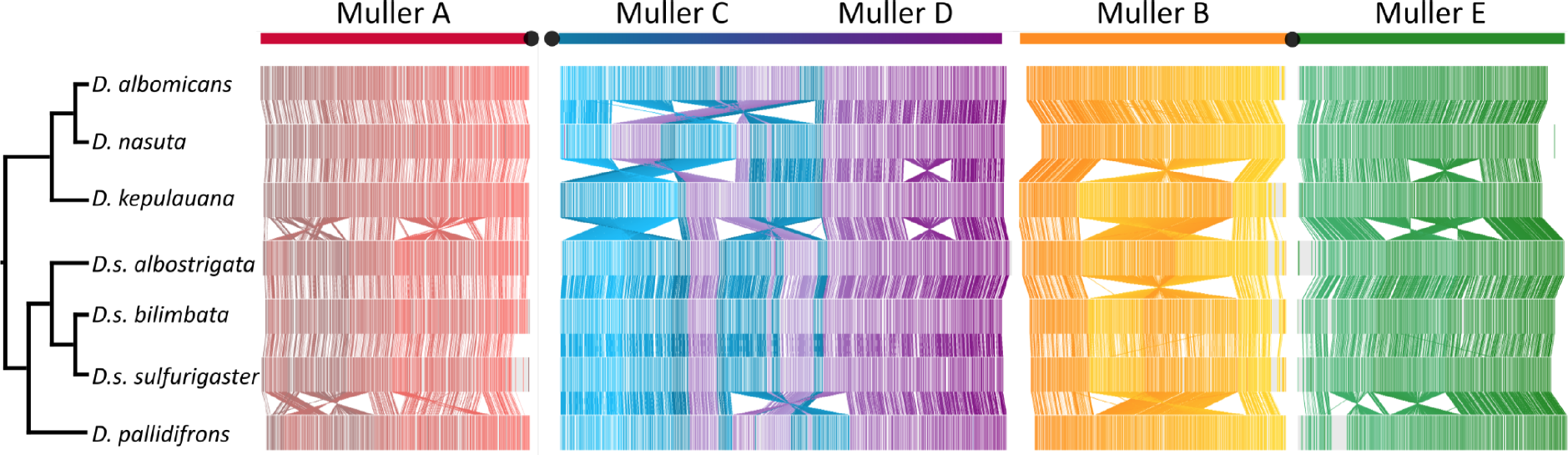
Phylogenetic relationship between species investigated and chromosomal synteny based on alignments of orthologous single-copy genes. Genes are color-coded according to their assignments to Muller elements. Note that Muller A and C/D are fused in *D. albomicans*.

### Identification of inversions using whole-genome alignments

We used whole-chromosome alignments to identify inversions on each chromosome arm for species of the *D. nasuta* subgroup. We used MUMmer to compare the chromosomes of each species and used breaks in synteny to map inversion breakpoints (Kurtz et al. 2004; **Figure 2**). For each chromosome arm, we identified syntenic segments and we used GRIMM to find the minimum number of rearrangements required to account for the order and orientation of syntenic segments along the phylogeny (Tesler 2002). In total, we identify 22 large chromosomal inversions along the major chromosomes (**Figure 3**). We identify 8 inversions on the ancestral X chromosome (Muller A), 6 inversions on the metacentric chromosome 2 (3 on Muller B and 3 on Muller E), and 8 inversion on the telocentric chromosome 3 (Muller C and Muller D). Therefore, while encompassing only a single Muller element and thus being substantially smaller than other chromosomes, the X has a similar number of inversions. Higher rates of X-linked inversions have also been found in primates (Porubsky et al. 2020). The chromosomal inversions identified vary dramatically in size, ranging from 3.9-18.0 Mb, and contain hundreds or thousands of genes (**Table 1**).

**Figure 2.**
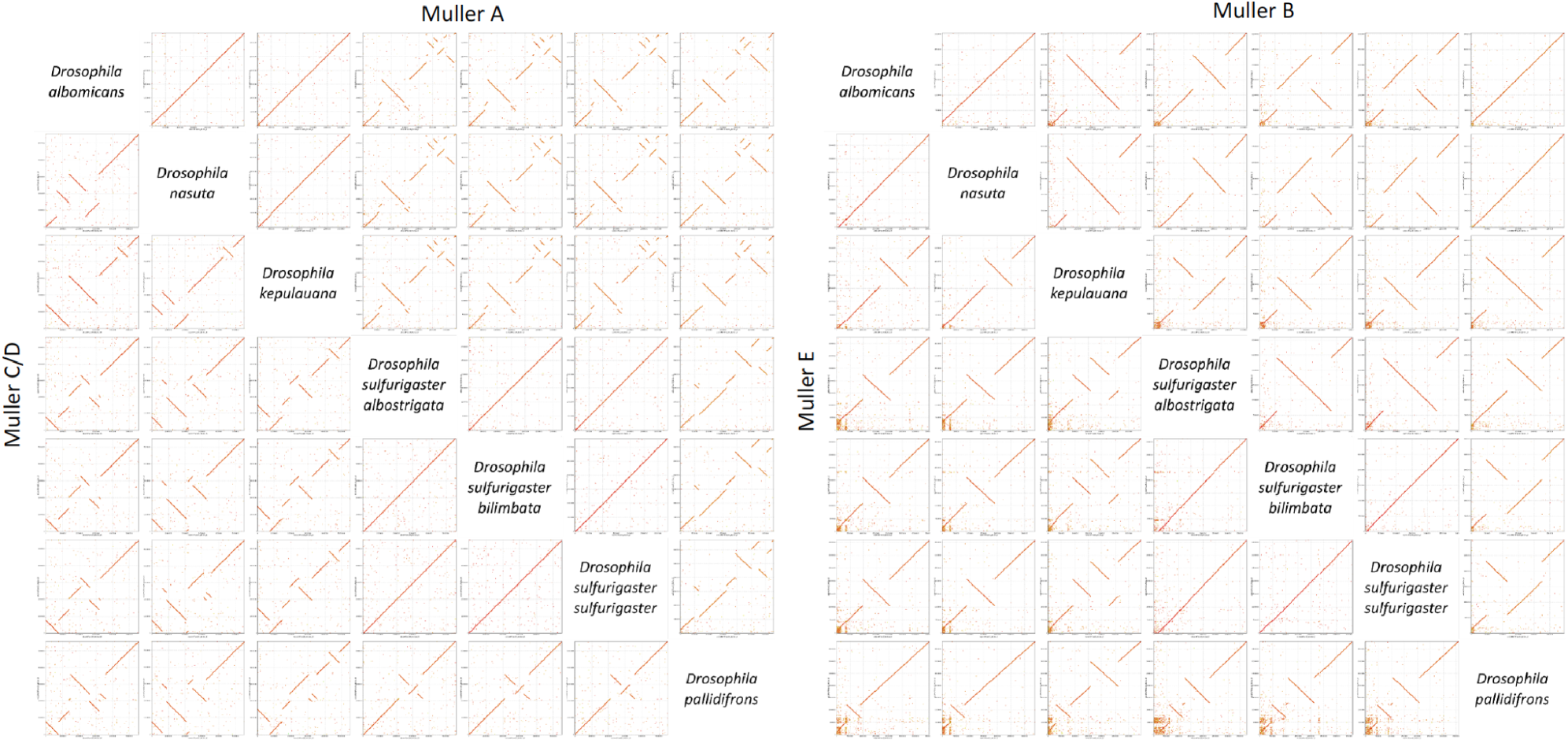
Dotplot between species for all major chromosome arms (Muller elements A through E). The pericentromere of each chromosome arm is placed at the bottom left corner of each subplot. Note that chromosomes are not drawn to scale (i.e. Muller C/D is approximately twice as large as all other chromosome arms).

**Figure 3.**
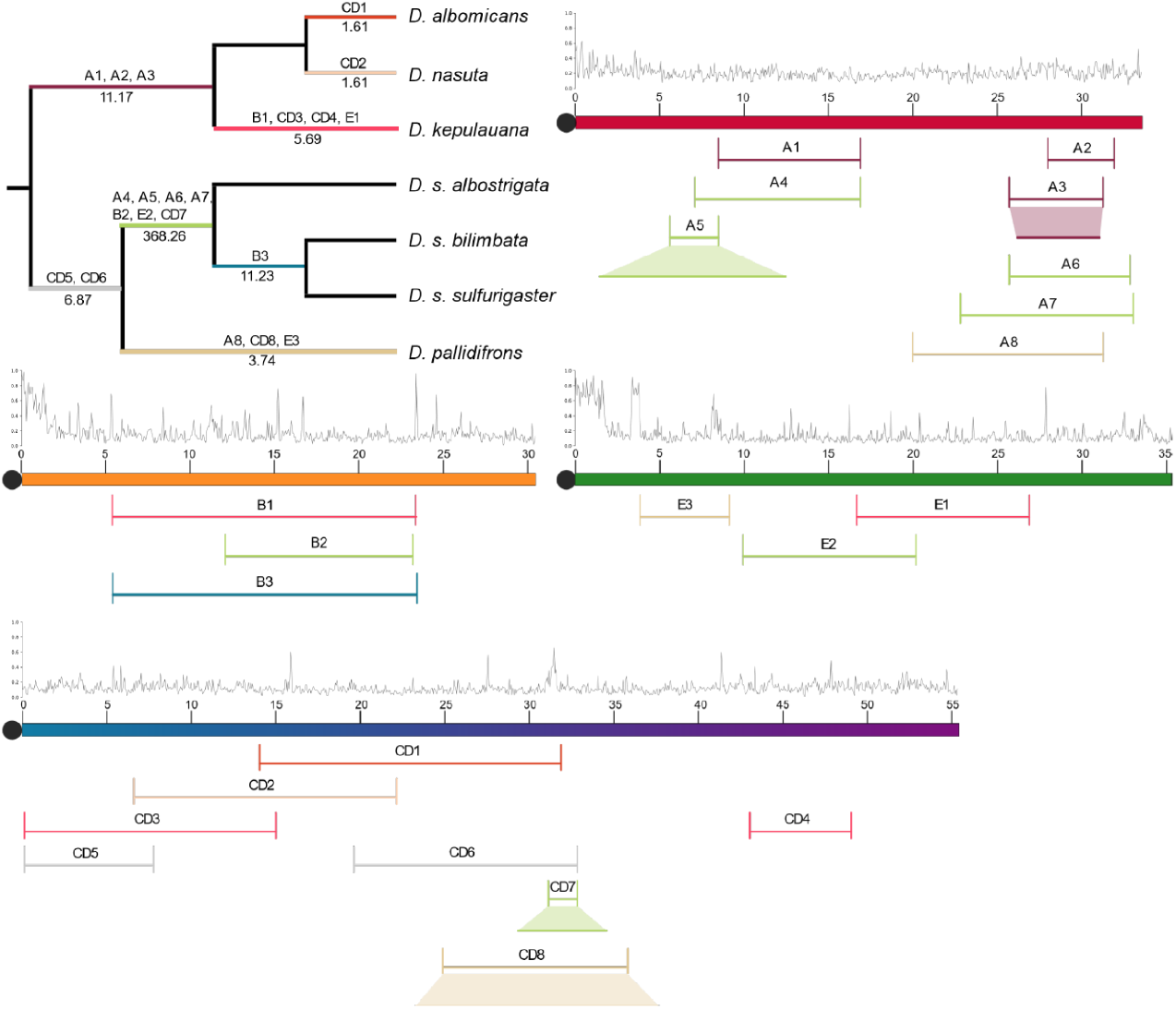
Major inversions along the phylogeny based on parsimony reconstruction. The approximate location of inversions along the ancestral chromosomes is indicated, and their size (the size of nested inversions is indicated under the shaded region). Inversions are color coded by the branch on the phylogeny where they occurred. Dots indicate location of the centromere and ticks mark every 5 Mb. The repeat content (fraction repeat masked in 50-kb windows) is shown above each chromosome. The number of inversions per million years is shown along each branch (bottom) of the phylogeny.

**Table 1.** Inversion sizes along the *nasuta* phylogeny

| Inversion | Size | Genome Proportion | Chromosome Proportion |
| --- | --- | --- | --- |
| A1 | 8,435,043 | 0.050 | 0.251 |
| A2 | 3,942,469 | 0.024 | 0.117 |
| A3 | 5,566,094 | 0.033 | 0.166 |
| A4 | 9,812,392 | 0.059 | 0.292 |
| A5 | 2,884,156 | 0.017 | 0.086 |
| A6 | 7,149,585 | 0.043 | 0.213 |
| A7 | 10,244,303 | 0.061 | 0.305 |
| A8 | 11,286,451 | 0.067 | 0.336 |
| B1 | 17,991,444 | 0.107 | 0.590 |
| B2 | 11,126,246 | 0.066 | 0.365 |
| B3 | 18,041,703 | 0.108 | 0.592 |
| CD1 | 17,855,989 | 0.107 | 0.322 |
| CD2 | 15,570,471 | 0.093 | 0.281 |
| CD3 | 14,885,804 | 0.089 | 0.268 |
| CD4 | 6,012,219 | 0.036 | 0.108 |
| CD5 | 7,631,642 | 0.046 | 0.138 |
| CD6 | 13,246,961 | 0.079 | 0.239 |
| CD7 | 1,720,990 | 0.010 | 0.031 |
| CD8 | 10,943,708 | 0.065 | 0.197 |
| E1 | 10,243,449 | 0.061 | 0.290 |
| E2 | 10,255,425 | 0.061 | 0.291 |
| E3 | 5,293,609 | 0.032 | 0.150 |

### Phylogenetic reconstruction of inversion

We reconstructed the evolution of inversions in the *nasuta* clade along the phylogeny using parsimony. **Figure 3** shows the inferred occurrence of inversions along different branches. Our sequenced strains of *D. albomicans* and *D. nasuta* differ by two overlapping inversions on Muller C/D (which forms the neo-sex chromosome in *D. albomicans*), but are otherwise co-linear. Their sister species *D. kepulauana* harbors two additional inversions on Muller C/D, and one on Muller B and Muller E, and this entire clade shares three inversions on Muller A. The sister species *D. s. sulfurigaster* and *D. s. bilimbata* are entirely collinear, and a single shared inversion on Muller B distinguishes them from their sister clade *D. s. albostrigata*. The *sulfurigaster* clade has four inversions on Muller A in common, and on each on Muller B, E and C/D. Their sister species *D. pallidifrons* has one inversion on Muller A, CD and E, and two inversions on Muller C/D occurred in the common ancestor of the *sulfurigaster* flies and *D. pallidifrons*.

Overall, we find the average inversion rate to be 5.2 inversions per million years, consistent with previously found inversion rates in *Drosophila* (Ranz et al. 2007; Lemeunier and Ashburner 1984; Powell 1997; Vieira et al. 1997; Bartolomé and Charlesworth 2006; Papaceit, Aguadé, and Segarra 2006; González, Casals, and Ruiz 2007; Bhutkar et al. 2008). However, there is high variation in the inversion rate per branch on the phylogeny (**Figure 3**). In particular, almost 1/3 of all inversions were identified on the short branch leading to species of the *sulfurigaster* species group.

In addition, inversions appear to be more common on the X chromosome compared to autosomes. While encompassing only about 1/5 of the total genome size, the X chromosome harbors more than 1/3 of all the inversions detected (**Figure 3**). Again, higher rates of inversions on the X chromosome are consistent with previous observations in *Drosophila* (Cheng and Kirkpatrick 2019).

### Molecular characterization of breakpoints

Localizing the precise inversion breakpoints can be informative for several reasons. Inversions may directly impact gene structure or gene expression, and the identification of inversion breakpoints might provide insights into the molecular mechanisms by which inversions arise. We therefore carefully characterized all the inversion breakpoints on Muller B and Muller E.

We identify three inversions on Muller B (**Figure 4**). Inversion B1 (which occurred along the *D. kepulauana* branch) and B3 (occurring along the *D. s. bilimbata*/*D. s. sulfurigaster* branch) are about 18 Mb in size. B1 and B3 occurred at homologous positions in the genome, and both of their breakpoints are located within the histone gene cluster. Nonalleleic homologous recombination could promote recurrent generation of inversions at the histone cluster, but it is also possible that this inversion was inherited from a common ancestor. Inversion B2 is about 11Mb long and shared by all *sulfurigaster* flies. One breakpoint of this inversion is located next to HP1 (*Su(var)205*), an important structural component of heterochromatin, but no repeated sequences are found at the breakpoints of the inverted chromosome (**Figure 4**).

**Figure 4.**
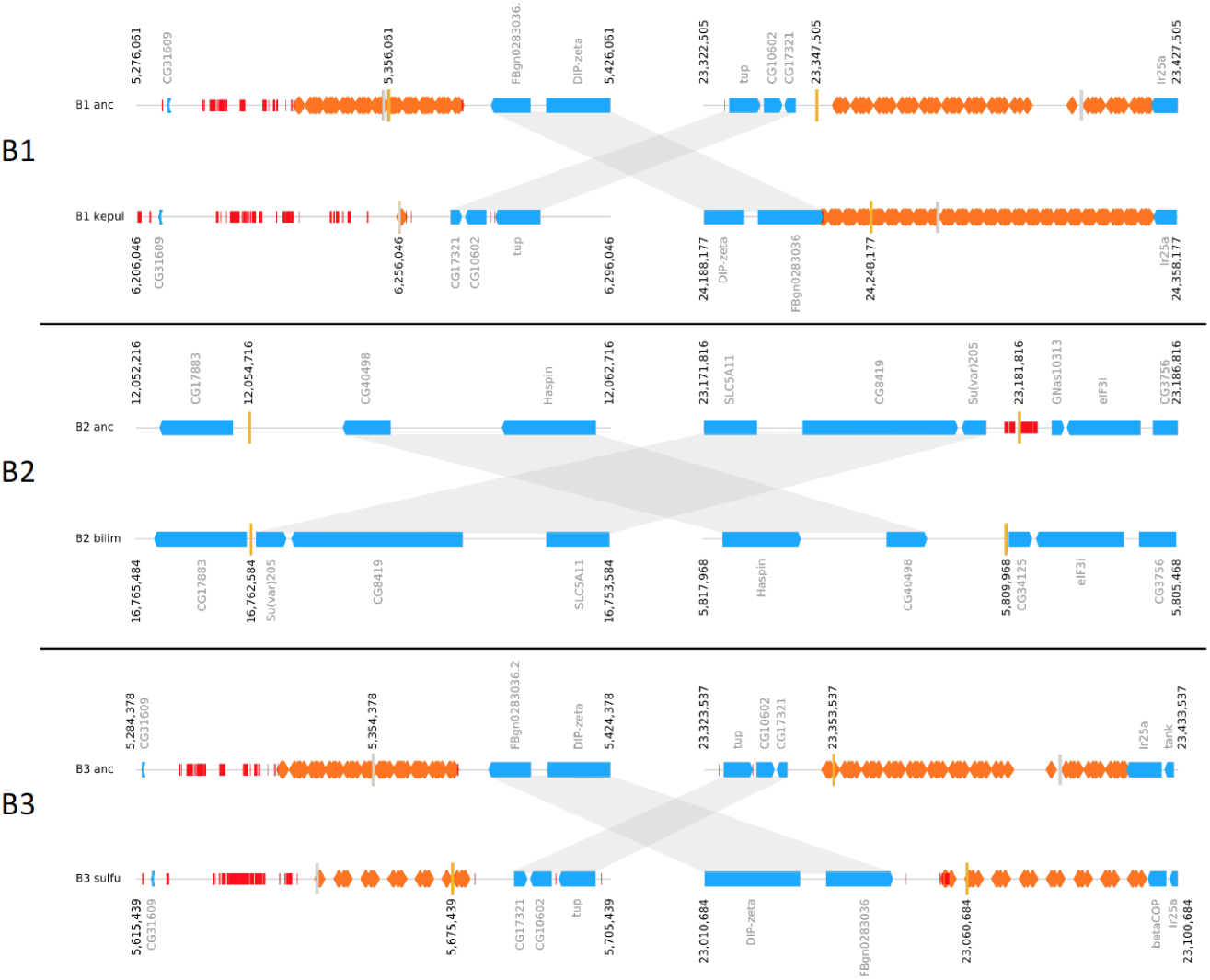
Inversion breakpoints on Muller B. Blue boxes indicate protein-coding genes, orange boxes indicate histone genes, and red boxes indicate repeats. Approximate breakpoint coordinates are given (yellow line), and homologous regions inside the inversion breakpoints are shown by grey shading.

Muller E harbors three inversions (**Figure 5**). Inversion E1 is about 10 Mb in size and occurred along the lineage leading to *D. kepulauana*. One of the breakpoints occurred at an approximately 1.6 kb repeat-masked region with no known homology aside from a 64 bp stretch that is homologous to R1-3_DF—a non-LTR retrotransposon. The other breakpoint is in a unique region (**Figure 5**).

**Figure 5.**
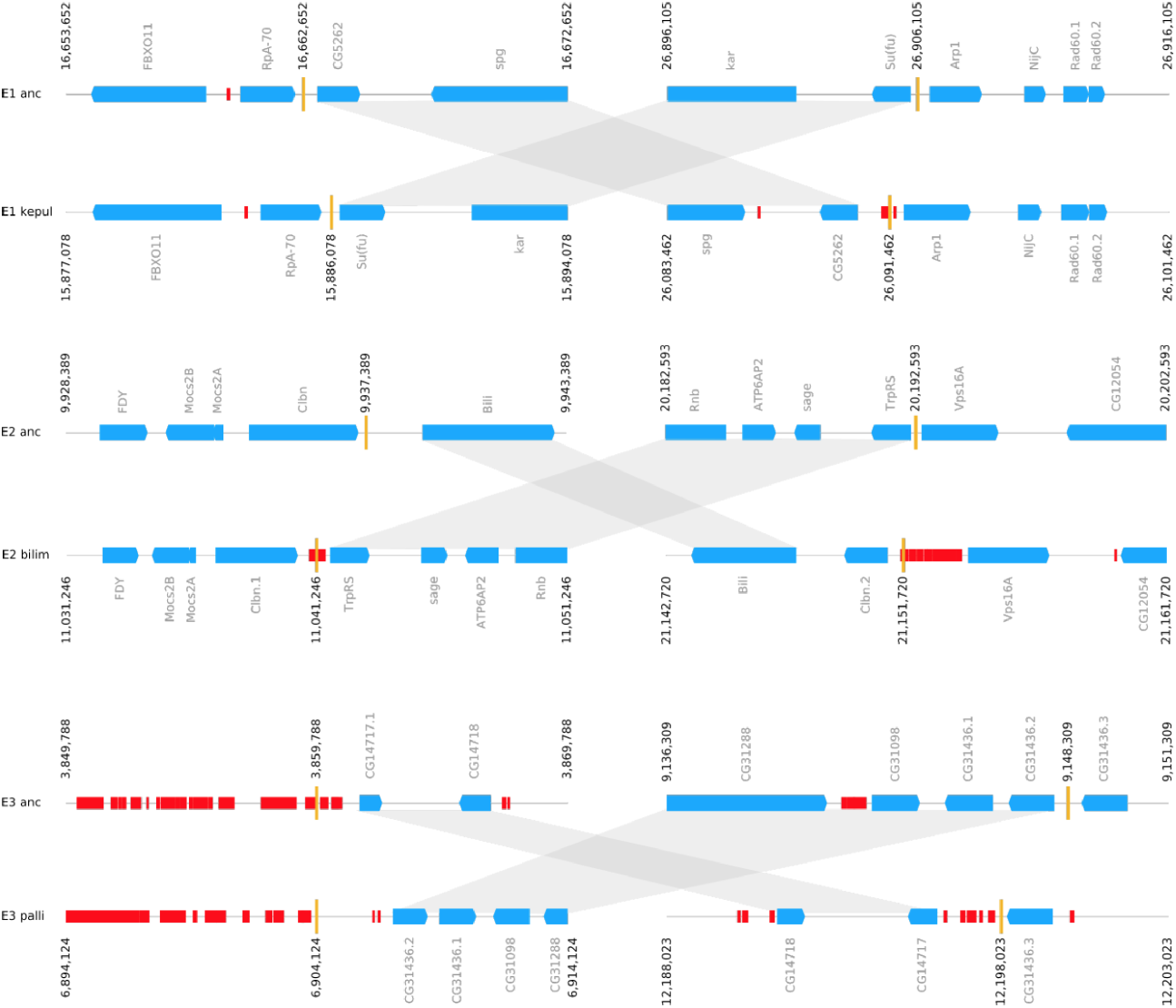
Inversion breakpoints on Muller E. For legend, see Fig. 4

Inversion E2 is about 10 Mb in size and occurred in the *sulfurigaster* lineage shared by *D. s. albostrigata, D. s. bilibmata,* and *D. s. sulfurigaster*. Both inversion breakpoints lie inside repetitive regions that are 1.2 kb and 13 kb in size (**Figure 5**). The breakpoint has a duplication of the *Clbn* gene along the *sulfurigaster* lineage while all other species in the *nasuta* clade have only a single copy of *Clbn*, suggesting that the inversion created a duplicate copy of this gene. Duplications of non-repetitive DNA at inversion breakpoints can be caused by staggered single-strand DNA breaks and repair by non-homologous end-joining (Guillén and Ruiz 2012). Inversion E3 occurred on the lineage leading to *D. pallidifrons* and is about 5 Mb in size. One inversion breakpoint is found inside a large (over 1 Mb long) repeat island, which in the *D. pallidifrons* genome is comprised of a number of transposons, over half of which are hAT elements. The other breakpoint is located within the tandemly duplicated multicopy gene *CG31436*. Thus, repeated sequences are found recurrently at inversion breakpoints in the *D. nasuta* species group.

## Discussion

Inversion polymorphism has been studied for over a century in *Drosophila*. Inversions can have profound biological influences (see introduction), but the evolutionary processes maintaining inversions are typically poorly understood. Individuals heterozygote for inversions may suffer reduced fertility by producing nonfunctional gametes during meiosis. These fertility effects are expected to be less pronounced in Drosophila, since males generally lack meiosis, and aberrant recombinant products contribute preferentially to the polar body nurse cells in females (Reis et al. 2018).

Large differences in rearrangement rates have been reported between species and between chromosomes in Drosophila. We find dramatic variation in the rate of chromosomal inversions among lineages. Seven out of the 22 inversions identified map to the short branch that leads to flies of the *sulfurigaster* species complex, but only a single inversion on the branch leading to *D. albomicans* or *D. nasuta* (**Figure 3**). This is in agreement with previous observations in *Drosophila*, which found that rates of chromosomal inversions can differ by over an order of magnitude even among closely related species and between Muller’s elements (Ranz et al. 2007; Lemeunier and Ashburner 1984; Powell 1997; Vieira et al. 1997; Bartolomé and Charlesworth 2006; Papaceit, Aguadé, and Segarra 2006; González, Casals, and Ruiz 2007; Bhutkar et al. 2008). This asymmetry in rates of inversions could result from differences in fitness effects or the efficacy of selection to establish new inversions, or from differences in mutation rates among lineages.

The molecular mechanisms of how inversions are generated are incompletely understood, and may differ among species or chromosomes. Inversions can be generated by nonallelic homologous recombination between repeated sequences, or by chromosome breakage and erroneous repair of the break by nonhomologous end-joining (Sonoda et al. 2006). Most inversions breakpoints in the *melanogaster* subgroup are associated with inverted duplication of genes or other non-repetitive sequences (Ranz et al. 2007). The presence of inverted duplications associated with inversion breakpoint regions was suggested to result from staggered breaks, followed by non-homologous end-joining. On the other hand, several studies in the *Drosophila* subgroup have found that repetitive elements are associated with the formation of inversion, suggesting an important role of ectopic exchange (Cacares et al. 1999; Fonseca et al. 2012). In the *D. nasuta* subgroup, we find evidence for both processes.

Reuse of inversion breakpoints in Drosophila has been reported at both the cytological and molecular level (Dobzhansky and Socolov 1939; Krivshenko 1963; Coluzzi et al. 1979; Lemeunier and Ashburner 1984; Pevzner and Tesler 2003; Zhao et al. 2004; Murphy et al. 2005; Richards et al. 2005; Goidts et al. 2005). We find that the histone gene cluster, which is located on two separate regions on Muller B in flies of the *nasuta* subgroup was involved in the generation of inversions in two separate lineages (though we cannot rule out that this inversion was segregating in a common ancestor of this species group). This resembles findings in great apes, where a high rate of homoplasy of inversions was observed (Porubsky et al. 2020). Reuse of inversion breakpoints might be due to mutational bias if these regions are particularly prone to breakage, or driven by selection if a specific breakpoint position affects the intrinsic fitness of a new arrangement (McBroome et al. 2020). Mutations caused by inversion breakpoints may have diverse consequences, from gene disruptions to generation of new gene duplicates or transfer of regulatory sequences from one gene to another. We identify one instance of a gene duplication generated by an inversion on Muller E in the *sulfurigaster* clade.

Chromosomal inversions can maintain linkage among alleles that are favored by natural selection and inversions that are associated with complex polygenic phenotypes are known from a variety of taxa (Hoffmann and Rieseberg 2008). Species from the *nasuta* clade are recently diverged, but differ in various morphological and behavioral phenotypes (Kitagawa et al. 1982; Spieth 1969). It will be of great interest to address the role of chromosomal inversions in contributing to phenotypic differences and local adaptation.

## Methods

### Genome Assemblies & Annotations

We used chromosome-level assemblies for seven species of the *D. nasuta* species group, which are described elsewhere (Wei, Mai et al, in preparation). **Table S1** lists the strains that were investigated. All assemblies are highly contiguous (N50s ranging from 34 Mb to 38 Mb) and very complete (with BUSCO scores ranging from 98.5% to 99.7%), and total assembly sizes ranging from 161 Mb to 163 Mb. For each species, the euchromatic portion of all Muller elements are assembled as a single contig, and only highly repetitive pericentromeric fragments could not be placed on the assembly (the assemblies comprise between 77 to 282 scaffolds with a mean of 157 scaffolds).

Gene annotations for each species (from Wei, Mai et al, in preparation) were clustered with *D. virilis* gene annotations using OrthoDB (Kriventseva et al. 2019). We then assign a name to each gene based on clustering with *D. virilis* annotations and their homology to *D. melanogaster* genes. An average of 10,534 genes were assigned to a *D. melanogaster* gene; 10,336 of these are single copy genes and 198 are duplicated genes.

### Inversion Along Phylogeny

MUMmer was used to determine the inversion status between all genome assemblies using the *D. albomicans* assembly as the reference (Kurtz et al. 2004). Sequences between each inversion breakpoint are assigned a numeric identifier and an optional negative sign to denote an inverted status relative to *D. albomicans* for each genome, which are then represented by an ordered sequence of these identifiers. The numeric sequence for each genome is then used as input for GRIMM to determine the optimal rearrangement scenario along a phylogeny, and identify the most likely ancestral genome structure (Tesler 2002).

We generated an “ancestral genome” by using the *D. albomicans* genome assembly, and ‘un-inverting’ all the inversions occurring along the branches leading to *D. albomicans* (that is, inversion A1, A2, A3, CD1; see Figure 3). To call inversion breakpoints, we took the mean between the end of the alignment on one side of the inversion and the start of the alignment on the other side of the alignment from MUMmer coordinates (**Table S2**).

All genome assemblies were then aligned to the ancestral genome using MUMmer. For each chromosome, all different inversion breakpoints are used to demarcate regions along the chromosome and inversions along the genome were then estimated using GRIMM based on the order and orientation of these regions relative to the ancestral genome.

### Inversion Rates

To test the rates of inversions along the phylogeny, we also calculate the branch lengths along the phylogeny. They are determined using the same method as Mai *et al.* 2019, using mean Ks values between species, a neutral mutation rate estimate of 3.46×10^−9^ per base per generation, and a 7 generation per year estimate for Drosophila (Zhang et al. 2006; Cutter 2008; Keightley et al. 2009). Internal branch lengths are calculated by subtracting the divergence time between sister species from the divergence time between the mean of the sister species and a outgroup. For example, let *d_AB_* be the divergence time between species A and B. Given the phylogeny ((A, B), C), the length of the branch from the root node to the shared node between A and B is calculated by 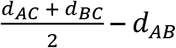.

The overall inversion rate is calculated using the total number of inversions on the phylogeny divided by the total length, in millions of years, of the phylogeny (in other words, the sum of all branches along the phylogeny). The inversion rate per branch is calculated by dividing the number of inversions occurring on the branch by the branch length.

## Acknowledgements

We acknowledge Ryan Bracewell for discussion.

---

**Table S1.** Strains investigated.

| species | strain | collection location |
| --- | --- | --- |
| *D. albomicans* | 15112-1751.00 | Okinawa, Japan |
| *D. nasuta* | 15112-1781.00 | Mysore, India |
| *D. kepulauana* | 15112-1761.03 | Ulu Temburong, Borneo |
| *D. s. albostrigata* | 15112-1771.04 | Rizal, Phillipines |
| *D. s. bilimbata* | 15112-1821.10 | Guam |
| *D. s. sulfurigaster* | 15112-1831.01 | Kavieng, New Ireland |
| *D. pallidifrons* | PN175_E-19901 | Ponape Micronesia |

**Table S2.** Inferred approximate inversion breakpoints.

| Inversion | Start<br>coordinates | End<br>coordinates | Size (kb) |
| --- | --- | --- | --- |
| Muller_A | 0 | 33597023 | 33,597 |
| A1 | 8461279.5 | 16896322.5 | 8,435 |
| A2 | 27993544 | 31936013 | 3,942 |
| A3* | 25697206.5 | 31263300 | 4,758 |
| A4 | 7083931 | 16896322.5 | 9,812 |
| A5* | 5577123.5 | 8461279.5 | 10,772 |
| A6 | 25697206.5 | 32846791.5 | 7,150 |
| A7 | 22802680.5 | 33046983 | 10,244 |
| A8 | 19976849 | 31263300 | 11,286 |
| Muller_B | 0 | 30469903 | 30,470 |
| B1 | 5356061 | 23347504.5 | 17,991 |
| B2 | 12054629 | 23180875 | 11,126 |
| B3 | 12053716 | 23181816 | 11,128 |
| Muller_DC | 0 | 55495487 | 55,495 |
| CD1 | 13906973 | 31762961.5 | 17,856 |
| CD2 | 6445133 | 22015604 | 15,570 |
| CD3 | 0 | 14885804 | 14,886 |
| CD4 | 42929924 | 48942143 | 6,012 |
| CD5 | 0 | 7631642 | 7,632 |
| CD6 | 19474208 | 32721168.5 | 13,247 |
| CD7* | 31000073 | 32721063 | 5,278 |
| CD8* | 24751727 | 35695434.5 | 14,491 |
| Muller_E | 0 | 35291776 | 35,292 |
| E1 | 16662651 | 26906100 | 10,243 |
| E2 | 9937162.5 | 20192587.5 | 10,255 |
| E3 | 3859696.5 | 9148309 | 5,289 |

\* nested inversion

